# Perilaryngeal Functional Muscle Network in Patients with Vocal Hyperfunction - A Case Study

**DOI:** 10.1101/2023.01.10.523514

**Authors:** Rory O’Keeffe, Seyed Yahya Shirazi, Sarmad Mehrdad, Tyler Crosby, Aaron M. Johnson, S. Farokh Atashzar

## Abstract

Patients with both phonotraumatic and non-phonotraumatic dysphonia commonly present with vocal hyperfunction, defined as excessive perilaryngeal muscle activity and characterized by muscular pain and strain in the neck, increased vocal effort, and vocal fatigue. The inability to reliably measure vocal hyperfunction is a barrier to adequate evaluation and treatment of hyperfunctional voice disorders. We have recently demonstrated that the perilaryngeal functional muscle network can be a novel sensitive neurophysiological window to vocal performance in vocally healthy subjects. In this paper, for the first time, we evaluate the performance and symmetry of functional perilaryngeal muscle networks in three patients with voice disorders. Surface electromyography signals were recorded from twelve sensors (six on each side of the neck) using the wireless Trigno sEMG system (Delsys Inc., Natick, MA). Patient 1 was diagnosed with primary muscle tension dysphonia, Patient 2 was diagnosed with unilateral vocal fold paresis, and Patient 3 was diagnosed with age-related glottal insufficiency. This paper reports altered functional connectivity and asymmetric muscle network scan behavior in all three patients when compared with a cohort of eight healthy subjects. Our approach quantifies synergistic network activity to interrogate coordination of perilaryngeal and surrounding muscles during voicing and potential <u>discoordination</u> of the muscle network for dysphonic conditions. Asymmetry in muscle networks is proposed here as a biomarker for monitoring vocal hyperfunction.

## 1. Introduction

Vocal hyperfunction (VH) is defined as excessive perilaryngeal muscle activation during phonation [1] and is associated with some of the most prevalent voice disorders [2], [3]. VH is considered the primary manifestation of primary muscle tension dysphonia (pMTD) [4]–[6]. In addition, VH contributes to the formation of vocal fold nodules [7], [8] and often emerges as a compensatory behavior in response to global insufficiency (e.g. vocal fold paralysis and age-related vocal fold atrophy) [9], [10]. In spite of the prevalence of related syndromes, clinical assessment of vocal hyperfunction is conducted with mainly subjective methods, such as patient self-reporting, palpation of the perilaryngeal region, perceived vocal effort, and laryngovideostroboscopy [11], [12].

Classic non-invasive surface electromyography (sEMG) has been investigated as a potential measure for the perilaryngeal muscle activation during phonation [13]–[16]. Previous sEMG-based investigations in normal or disordered voice analyzed the temporal and spectral characteristics of the sEMG from one or two muscles [17]. The reliability of sEMG recording from perilaryngeal muscles is typically excellent [18]. However, studies using sEMG measures such as root mean square (RMS) and power spectral density (PSD) of single muscles have shown heterogeneous outcomes for assessment of VH, with some studies suggesting differences in sEMG parameters between patients with muscle tension dysphonia and controls, [19], [20] while others are finding no differences at rest or in various phonatory tasks [21]. This heterogeneity is likely because voicing is a complex neuromuscular activity, and activation differences in one or two single muscles are unlikely to capture this complexity. There exist studies designed to address this limitation by assessing beta-band (15-35 Hz) coherence between two anterior neck muscles during voicing, which showed some discriminative power to indicate hyperfunction and differences between control subjects and patients with vocal nodules [22], [23]. Other recent studies show the promise of multichannel sEMG to distinguish vocal dysfunction from typical voicing [15], [24]. In this case study, we investigate the potential of the functional muscle network as a biomarker of VH.

The functional muscle network is an emerging concept that uses simultaneous multichannel sEMG to decode muscle synergy during complex motor tasks [25], [26]. In our recently published study, we used multichannel sEMG to expand from a single coherence measurement in specific frequency bands to a wideband intermuscular coherence network, thereby increasing the possibility for quantifying the complex neuromuscular activity of perilaryngeal muscles during voicing due to wider spectral and spatial distribution of the analysis [27]. Our approach showed different muscle network features discriminated different vocal tasks in typical (non-disordered) speakers.

In this paper, we evaluate the performance of functional perilaryngeal muscle network connectivity during voicing using synchronized multi-sensor sEMG. The level of coordination in perilarygneal muscles during voicing and the level of <u>discoordination</u> during dysphonic voicing is estimated by investigating the intermuscular coherence network, which measures the underlying spectrotemporal synergies. As mentioned, we have recently and for the first time, proposed the concept of perilaryngeal muscle network across different vocal tasks in normal speakers (e.g., sustained vowels, pitch glide), [27], and we showed aberrant network activity in hyper-functional speakers. In this paper, we take the next step and evaluate the performance of functional muscle networks in three cases of vocal hyperfunction with three different voice disorder diagnoses. The paper suggests asymmetry in perilaryngeal functional muscle connectivity as a biomarker of VH.

The rest of this paper is organized as follows. In Section II, we will provide the method of data collection and network formulation and will explain the objective metrics quantifying the integration and segregation of neural information at the muscle networks. In Section III, we provide the results comparing muscle networks in VH patients with that of the healthy cohort. Concluding remarks are given in Section IV.

## II. Method

The institutional review board of the New York University Grossman School of Medicine approved the study. Three female patients with a voice disorder diagnosed by a fellowship-trained laryngologist and symptoms of VH were recruited at the NYU Langone Voice Center. Patient 1 (35 years old) was diagnosed with primary muscle tension dysphonia, Patient 2 (56 years old) was diagnosed with unilateral right vocal fold paresis, and Patient 3 (57 years old) was diagnosed with age-related glottal insufficiency and bilateral sulcus. All patients presented with symptoms of vocal hyperfunction (e.g. increased effort with voicing, perilaryngeal muscle soreness or fatigue with voicing). Also, data were collected from eight subjects (four females) without a known history of voice disorders. After providing written informed consent, subjects performed a series of vocal tasks with varying degrees of pitch and loudness.

Three groups of tasks were tested. The first group of tasks involved sustaining the vowel /a/ at a constant pitch and volume, at four combinations of two levels of loudness (habitual/increased) and two levels of pitch (habitual/high pitch). Subjects performed three trials of each task. The second group of tasks included single repetition vocal exercises, namely (i) pitch glide [28], (ii) spontaneous speech, and (iii) singing. The third group of tasks involved reading the first paragraph of The Rainbow Passage, a reading commonly used in voice and speech research, at three levels of loudness: whispering, normal, and elevated [29].

sEMG signals were recorded from twelve sensors (six on each side of the neck) placed parallel to the direction of the muscles using the wireless Trigno sEMG system (Delsys Inc., Natick, MA) with a sampling frequency of 1259 Hz (Fig. 1). Four bipolar Trigno Mini sensors were used for the smaller perilaryngeal muscles (inferior and superior infrahyoid, bilaterally), while eight bipolar Trigno Avanti sensors were used for masseter, superior sternocleidomastoid, inferior sternocleidomastoid, and trapezius. Signals were processed using MATLAB R2020b (MathWorks Inc. Natick MA). The signals were filtered with a high-pass filter at 20

**Fig. 1.**
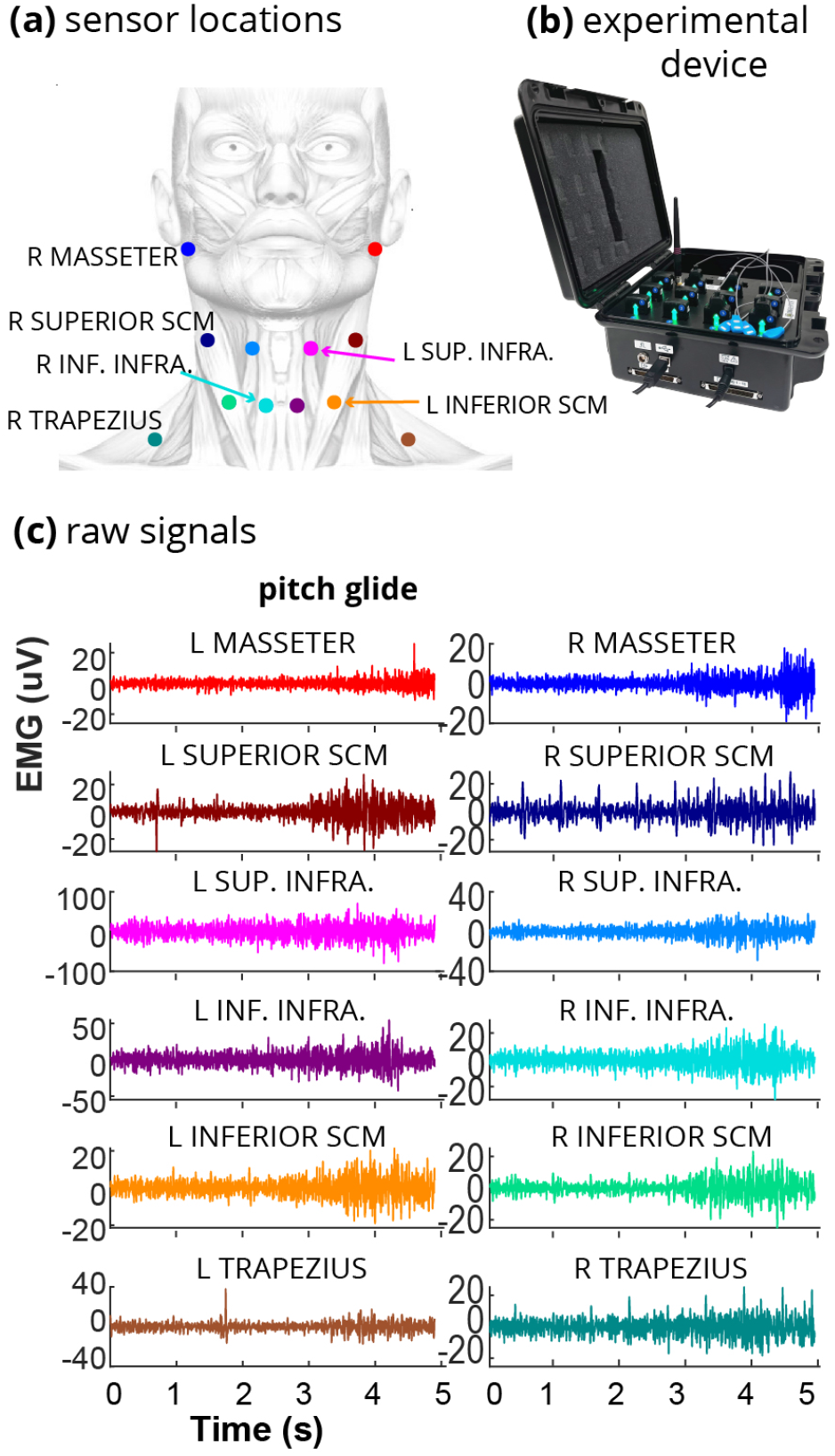
**(a)** Six sensors were placed at the masseter, superior sternocleidomastoid, superior infrahyoid, inferior infrahyoid, inferior sternocleidomastoid, and trapezius muscles. **(b)** sEMG was recorded using the wireless Trigno system (Delsys Inc., Natick, MA) Avanti and Mini sensors (blue head). **(c)** An exemplar recording from all muscles indicates modulations of the sEMG during the pitch glide.

Hz, a band-stop filter at 57.5-62.5 Hz for power-line noise, and a low-pass filter at 100 Hz. Thus, we considered the 20-100 Hz range. An exemplar recording from all muscles (indicating modulations of the sEMG during the pitch glide) is shown in Fig. 1(c).

### A. sEMG Network Analysis

Muscle networks were constructed for all tasks, using magnitude coherence analysis that estimates the frequency-based power transfer between two signals and has been used in a wide range of neuroscientific and neurophysiological studies, mainly at the central nervous system level, to decode information sharing between two neural activities. Magnitude squared coherence, *C*_*xy*_ between two signals *x*(*t*) and *y*(*t*) is:

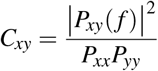

where *P*_*xx*_ and *P*_*yy*_ are the power spectral densities (PSDs) and *P*_*xy*_ is the cross power spectral density (CPSD).

To compute the coherence, Welch’s overlapped averaged periodogram method was utilized with a Hamming window of 2048 samples (1.63 ms) and 50% overlap. [30] The median coherence component in the 20-100 Hz range was considered for each connection edge between two sensor pairs to finally generate the proposed muscle networks consisting of multiple edges and nodes (muscles) for each trial. The network can be visualized using the net shape or adjacency matrix. In the case of tasks that had multiple trials, the median network across trials was computed in this work. In order to objectively quantify the topographical changes in muscle networks, we used measures from network sciences, including network degree, clustering coefficient, and global efficacy. The degree of each node, *D*_*i*_, is the average of all edges connected to the node and can be calculated as:

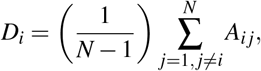

where *N* is the number of nodes, and *A*_*i j*_ is the *i j*^*th*^ element of the Adjacency matrix of the network. In this context, a node’s weighted clustering coefficient (*WCC*_*i*_) is defined as the measure of how well that node is connected to its local neighbors. The weighted clustering coefficient is defined as:

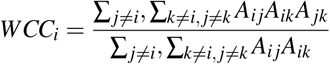

## III. Results

The healthy controls showed a significant increase in response to raised loudness and pitch (Figs. 2, 3). Compared to the healthy cohort, patients showed heterogeneous pairwise muscle coherence indicated in network scan analysis (Figs. 3, 4). The adjacency muscle network of the three patients for three tasks of interest is shown in Fig. 4. Overall, (i) the network scans showed a lower mean degree for the patients compared to the healthy cohort (Fig. 3), (ii) the network scans showed skewed and asymmetric response towards one side during certain tasks (Fig. 4), and (iii) the network scans showed lack of consistent modulations with pitch and loudness, in contrast to the monotonic response observed in the healthy cohort for the network mean degree (Fig. 3).

**Fig. 2.**
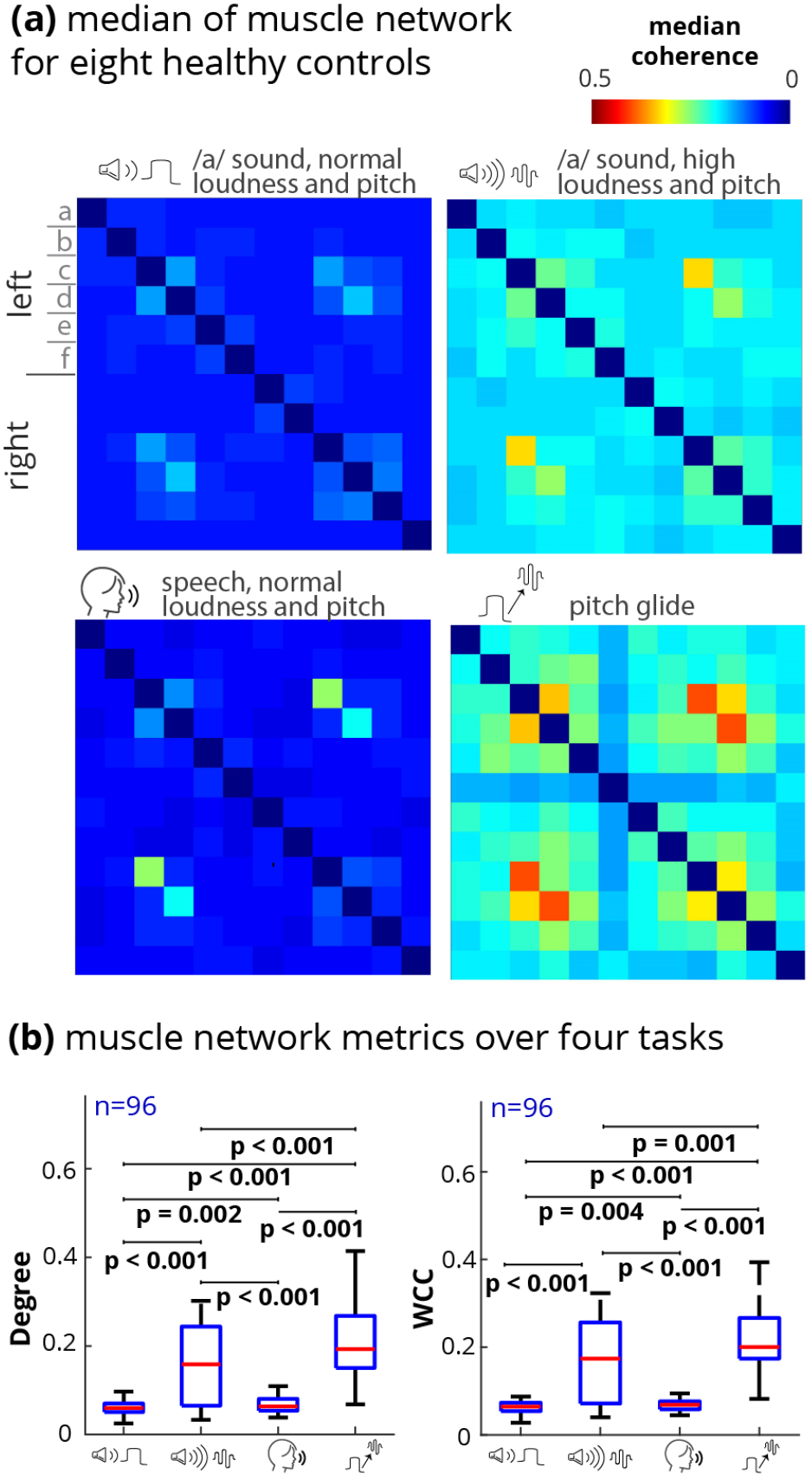
The results in this figure summarize our recently published findings [27]. It is presented here as the point of comparison for the evaluation of the muscle network in VH patients for the first time. **(a)** The median (over subjects) of adjacency matrices of functional muscle network from 8 healthy adults. The adjacency matrices are constructed from the pairwise median coherence between the sEMG signals during each task. The muscles are denoted as follows: a = masseter, b = superior sternocleidomastoid, c = superior infrahyoid, d = inferior infrahyoid, e = inferior sternocleidomastoid, f = trapezius. **(b)** The degree and weighted clustering coefficient of the network scans show a gradual increase with the increases in pitch and loudness.

**Fig. 3.**
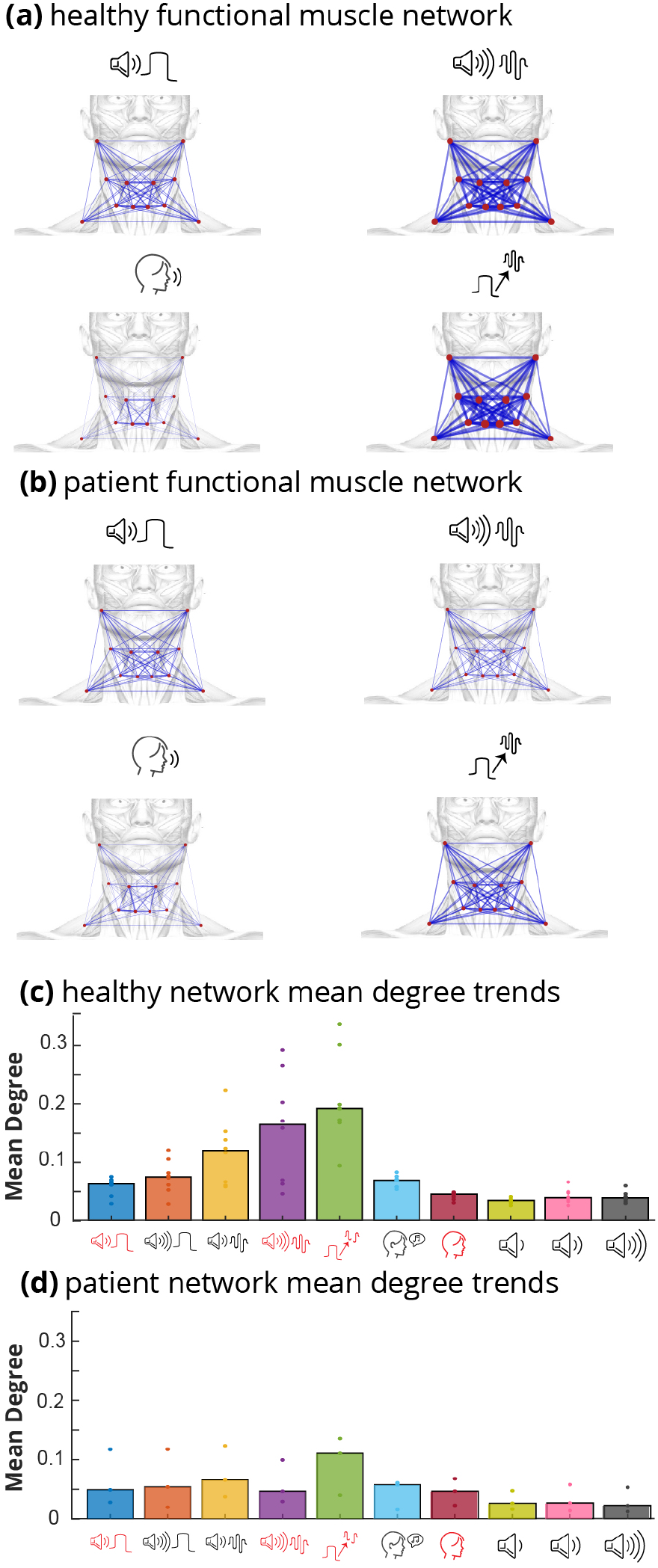
**(a)** Median of 8 functional muscle networks from 8 healthy adults [27]. **(b)** Median of 3 muscle networks from 3 patients. **(c)** Mean network degree for 8 healthy subjects over all vocal tasks. The network for the healthy cohort is symmetrical and strongly responds to task change. The ten vocal tasks are shown on the x-axis. The first four tasks are the sustained phonation tasks with (1) habitual loudness, habitual pitch, (2) elevated loudness, habitual pitch, (3) habitual loudness, high pitch, and (4) elevated loudness, high pitch. Tasks 5-7 are the single repetition tasks as follows: (5) pitch glide, singing, and (7) spontaneous speech. Tasks 8-10 are the reading tasks with (8) whispering loudness, (9) habitual loudness, and (10) elevated loudness. **(d)** Mean network degree for three patients over all tasks. Pitch glide has the highest mean degree for all subjects.

**Fig. 4.**
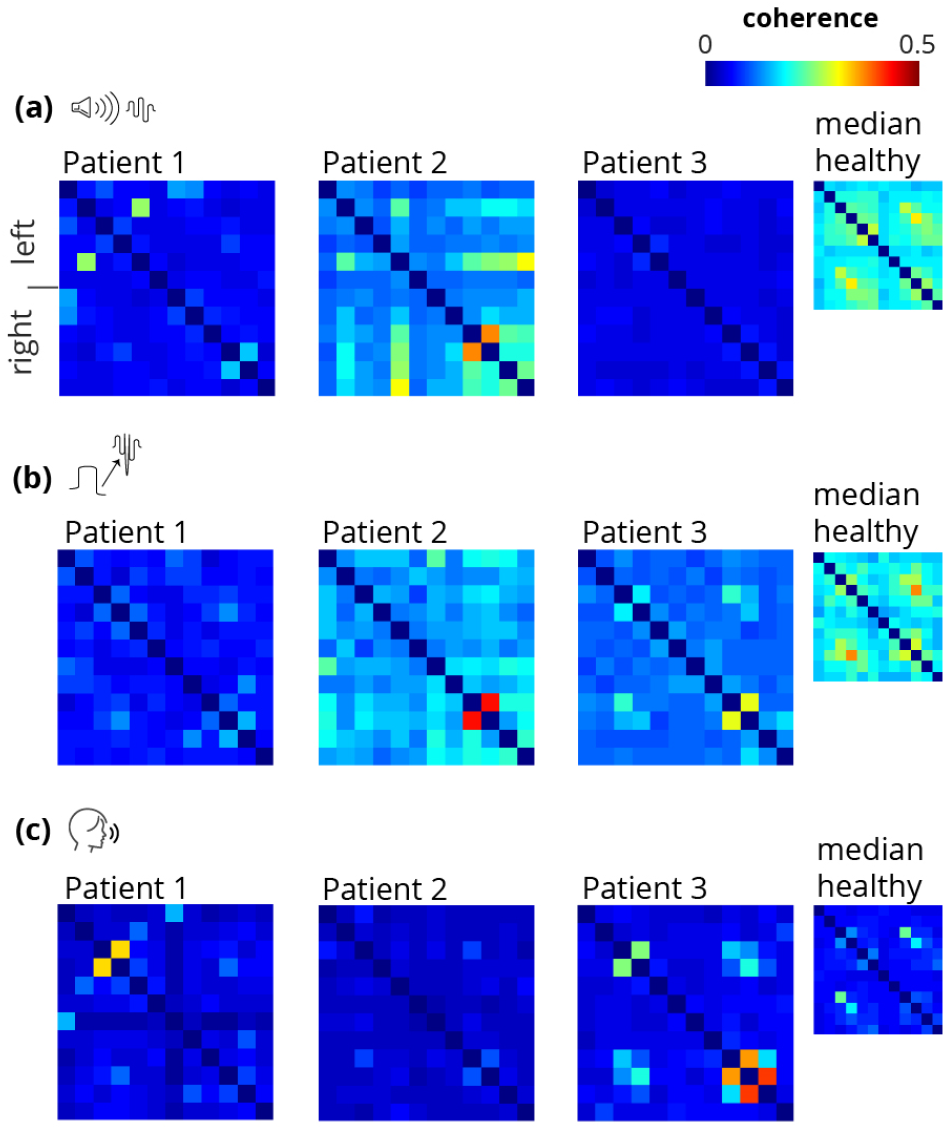
Adjacency matrices of three patients compared to the corresponding median adjacency matrix of the healthy cohort, for three tasks of interest. For the /a/ task with high loudness and pitch, Patient 2 indicated that connectivity was skewed towards the right side of the network (right-sided asymmetry). **(b)** For the pitch glide task, Patient 2 indicated right-sided asymmetry. **(c)** For the speech task, the network connectivity of Patient 1 was skewed to the left while Patient 3 showed a right-sided asymmetry.

### A. Sagittal symmetry is disturbed in dysphonic patients

To further analyze the network behavior in our preliminary cohort, the sagittal asymmetry matrix was computed by (i) splitting a muscle network adjacency matrix *A* into two sub-networks, left (*A*_*L*_) and right (*A*_*R*_), and (ii) computing the sagittal asymmetry matrix:

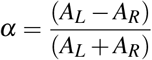

The sagittal asymmetry matrix *α* displays both the magnitude and direction of asymmetry. If the left-side asymmetry is positive, this implies that the left edge in the network scans dominates the corresponding right edge in terms of connectivity. Sagittal asymmetry provides a metric for quantifying the mediolateral changes of the coherence network and indicates how one side of perilaryngeal network scans dominates connection during functional voice tests.

After calculating *α* for a given task, the mean asymmetry across each muscle was calculated. The resultant 3-D muscle-task-asymmetry maps are illustrated in Fig. 5 for each patient, alongside the median map across all controls (Median Control). In the 3-D asymmetry map, the x-, y-, and z-axis respectively indicate the left-side muscles, the vocal tasks performed by the subjects, and the asymmetry value. Note that in this figure, a right-sided asymmetry will be seen as negative values. Patient 1 showed a left-sided dominance for the asymmetry (positive asymmetry), while Patients 2 and 3 showed right-sided dominance (negative values) (Fig. 5). As can be seen, each patient showed a unique and high sagittal asymmetry response, indicating that the muscle network is sensitive to personalized impairments, their severity, function, and sites.

**Fig. 5.**
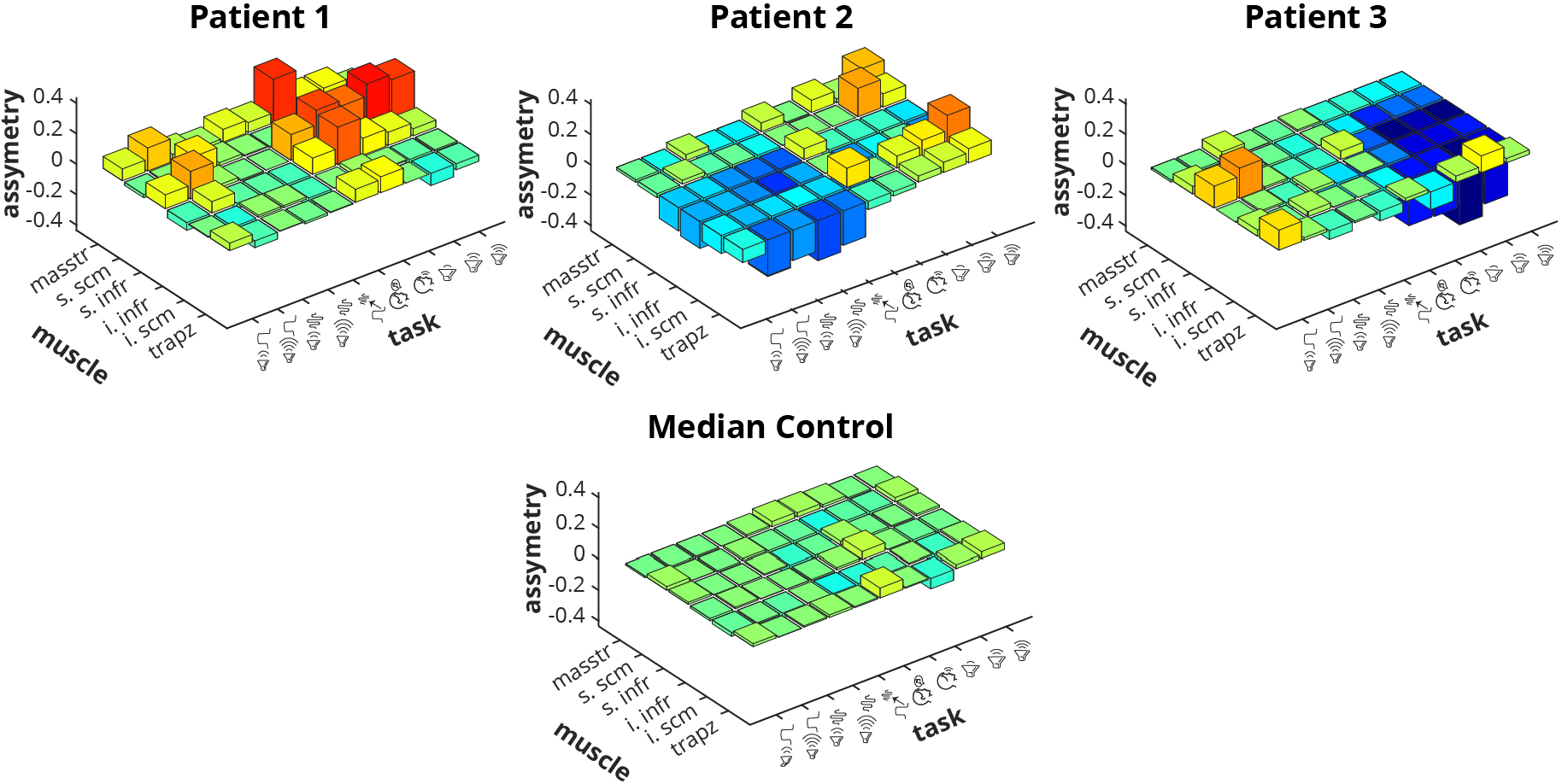
3D sagittal asymmetry maps show high magnitude of asymmetry (| asymmetry |) in patients, relative to controls. The x-, y-, and z-axis respectively indicate left-side muscles, vocal tasks performed, and asymmetry values. The ten vocal tasks are shown on the y-axis. The first four tasks are the sustained phonation tasks with (1) habitual loudness, habitual pitch, (2) elevated loudness, habitual pitch, (3) habitual loudness, high pitch, and (4) elevated loudness, high pitch. Tasks 5-7 are the single repetition tasks as follows: (5) pitch glide, (6) singing, and (7) spontaneous speech. Tasks 8-10 are the reading tasks with (8) whispering loudness, (9) habitual loudness, and (10) elevated loudness. Red indicates left-dominant asymmetry (positive score), and blue is a right-dominant asymmetry (negative score) in the network scan. Patient 1 showed strongly right-dominant while Patients 2 and 3 showed strongly left-dominant asymmetry for certain tasks.

In Fig. 6, the top 25% absolute values from the asymmetry maps of the patients are shown in comparison to the top 25% of the control cohort. The violin plot for each patient displays the absolute of the top 25% elements in each muscle-task | asymmetry | map (*n* = 0.25*×* #muscles*×* #tasks = 15). The violin plot for the ensembled healthy displays together the absolute of the top 25% elements from each control’s asymmetry map (*n* = 0.25*×* #muscles*×* #tasks*×* #subjects = 120). The absolute asymmetry violin plots show that the three patients had a high asymmetry when compared with the eight healthy subjects (*p <* 0.001), using Wilcoxon rank-sum test at the 5% significance level with Bonferroni correction. It is worth noting that the network modulations also differ per task across the patients. The asymmetry of Patients 1 and 3 is higher during the reading tasks, while the asymmetry of Patient 2 is more elevated during phonation (/a/ and pitch glide) tasks.

**Fig. 6.**
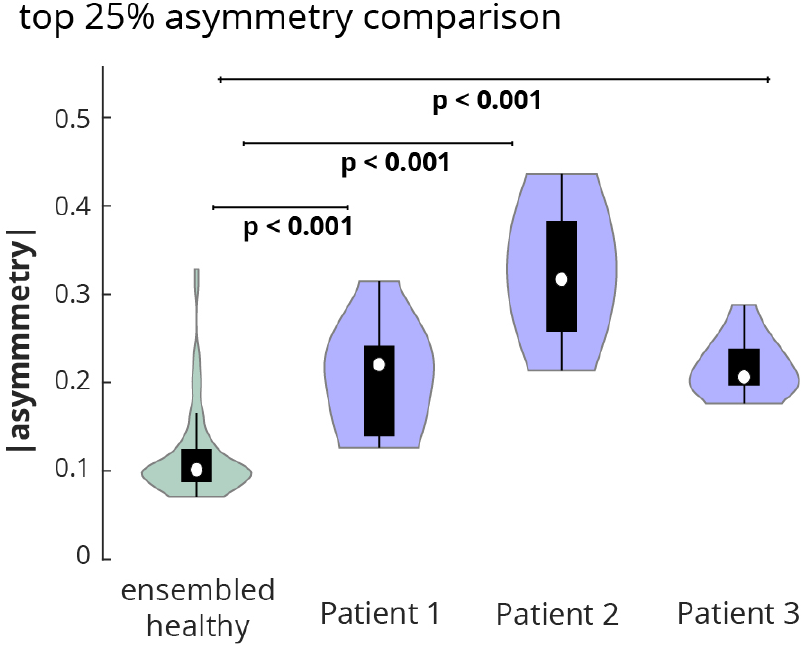
Each patient has higher top 25% of | asymmetry| than the healthy controls. The top 25% of| asymmetry| across the muscles and tasks are shown for each patient’s violin plot. The top 25% of | asymmetry | across the muscles and tasks were computed for each control and combined for the ensembled healthy violin plot. The null hypothesis was rejected for each comparison of the ensembled healthy with a patient (Wilcoxon rank-sum: *p <* 0.001).

## IV. Conclusion

In this paper, we proposed the concept of alternation in the perilaryngeal functional muscle network as a potential biomarker of vocal hyperfunction. For this, the study includes eight healthy subjects and three patients (one diagnosed with primary muscle tension dysphonia, one diagnosed with unilateral vocal fold paresis, and one diagnosed with age-related glottal insufficiency). Surface electromyography data were collected from 12 channels, and median magnitude coherence was considered as the connectivity method. The results support that all three patients showed asymmetric muscle networks when compared with the healthy cohort. At the same time, the type of asymmetry in each patient was condition-specific. It should also be noted that the results showed that the modulation of mean network degree in healthy subjects is more responsive to the increased pitch and loudness, while patients with vocal hyperfunction showed less responsive mean network degree for their perilaryngeal muscle network. The study was limited by the number of subjects.

